# The Exon Junction Complex component EIF4A3 plays a splicing-linked oncogenic role in Pancreatic Ductal Adenocarcinoma

**DOI:** 10.1101/2023.12.07.570542

**Authors:** Ricardo Blázquez-Encinas, Emilia Alors-Pérez, María Trinidad Moreno-Montilla, Víctor García-Vioque, Marina Esther Sánchez-Frías, Andrea Mafficini, Juan L. López-Cánovas, Corinne Bousquet, Manuel D. Gahete, Rita T. Lawlor, Raúl M. Luque, Aldo Scarpa, Álvaro Arjona‐Sánchez, Sergio Pedraza-Arevalo, Alejandro Ibáñez-Costa, Justo P. Castaño

## Abstract

Pancreatic ductal adenocarcinoma (PDAC) is one of the most lethal cancers worldwide and further research on its biology is needed to fully understand the disease. Dysregulation of alternative RNA splicing is a common hallmark in cancer, including PDAC, which provides an emerging source of knowledge and of novel biomarkers and therapeutic tools. Here, we examined the role of EIF4A3, a core component of the Exon Junction Complex intimately linked to RNA splicing, in pancreatic cancer biology. EIF4A3 is overexpressed in PDAC tissue and associated to clinical parameters of malignancy and poorer patient survival. Mechanistically, exploration of PDAC RNA-seq data unveiled the link of EIF4A3 to diverse malignancy processes, in line with its association to key molecular pathways. Accordingly, EIF4A3 targeting *in vitro* decreased essential functional tumor features such as proliferation, migration, colony formation and sphere formation, while its *in vivo* targeting reduced tumor growth. *EIF4A3* silencing in PDAC cell lines severely altered its transcriptional and spliceosomic landscapes, as shown by RNA-seq analyses, suggesting a role for EIF4A3 in maintaining RNA homeostasis. Our results indicate that EIF4A3 dysregulation in PDAC has a pleiotropic regulatory role on RNA biology, influencing key cellular functions. This paves the way to explore its potential as a novel biomarker and actionable target candidate for this lethal cancer.

## INTRODUCTION

Pancreatic ductal adenocarcinoma (PDAC) remains one of the most lethal cancers worldwide, mostly due to its late diagnosis —often with metastasis— and current lack of effective treatments, jointly leading to poor prognosis [1–3]. Research advances on PDAC molecular biology have established the primary pathologic relevance of the most frequently mutated genes: *KRAS, TP53, CDKN2A* and *SMAD4* [1, 2]. However, the knowledge gathered to date cannot fully explain the molecular underpinnings of the disease, and its present translation into clinical benefits for the patients is still lagging [2, 4–6]. Hence, novel research avenues are sorely required to find new molecular players that facilitate a better understanding of PDAC and provide new biomarkers and therapeutic targets to tackle this pathology. In this vein, studies from our group and others have shown that dysregulation of RNA splicing plays a key role in PDAC, which leverages this still limitedly known mechanism as a source of novel molecular tools [7–12].

Splicing is a process of RNA maturation that occurs co-transcriptionally, by which introns are removed from pre-RNA and exons are joined together [13]. This process is finely regulated to enable the generation of different mature RNA variants, or isoforms, from the same gene, which can then give rise to different proteins, or to mature non-translated RNAs [14], thereby increasing the diversity and versatility of the genome. Alternative splicing is therefore a capital step within the central dogma of Biology (DNA->RNA->protein) and its dysregulation is linked to the appearance of many diseases including cancer [15]. Altered splicing can lead to the generation of oncogenic splice variants and/or changes in signaling pathways, which contribute to cancer development, progression, aggressiveness, metastasis and drug resistance [16, 12].

RNA splicing is intimately intertwined with other processes involved in RNA biology, whose machineries often share molecular components and interact to ensure the correct maintenance of RNA processing and metabolism [17]. In this context, a paradigmatic mechanism is the Exon Junction Complex (EJC), a key interactive player that participates in, regulates, and interrelates several RNA-related processes such as splicing, RNA export from the nucleus, translation, m6A RNA methylation, and RNA degradation [18, 19]. A central protein of the EJC is EIF4A3, which together with MAGOH and RBM8A forms a trimeric core that interacts with many other proteins [18]. EIF4A3 is a DEAD box-family RNA helicase that shares high homology with other two genes from this family, *EIF4A1* and *EIF4A2*, which participate in translation initiation, and have been implicated in cancer [18, 20]. EIF4A3 does not participate in translation initiation, but its main function has long been related to the EJC [18], while an emerging number of additional functions, related to RNA biology, are being proposed for this molecule: from its early link to nonsense mediated decay (NMD) [21] to its recent bonds to ribosome biogenesis and m6A regulation [22], along with multiple associations with noncoding RNA regulation, including long noncoding RNA (lncRNA), circular RNA (circRNA), and micro RNAs (miRNA; [23], reviewed in [18]. Previous studies on the EIF4A family in PDAC mostly focused on EIF4A1/EIF4A2 and their role in translation initiation [24–26]. In contrast, few reports have examined the RNA metabolism-related functions of EIF4A3 in PDAC, where it was shown to mediate the actions of a lncRNA, LINC01232 [27] and a circRNA, circRNF13 [28]. Actually, the first identification of EIF4A3 in PDAC was unrelated to its currently known functions, but derived from a proteomic-based detection of Dead-box protein 48 (DDX48, an earlier name for EIF4A3) as an autoantigen in the sera of PDAC patients [29]. While it was proposed that the detection of autoantibodies to DDX48 could help improve PDAC diagnosis, no further reports confirmed or extended this idea. In other cancers, EIF4A3 has been found to be dysregulated and to play an oncogenic role and its underlying mechanisms have been examined [30–32]. Recently, we and others reported a profound alteration of key components of the splicing machinery in PDAC, with pathological implications [7–11]. This prompted us to devise the present study, aimed at elucidating the role of *EIF4A3* in PDAC pathophysiology, with particular attention to its involvement as a regulator of alternative splicing.

## MATERIALS AND METHODS

### Patients and samples

Formalin-fixed paraffin embedded (FFPE) samples from 75 PDAC patients were used in this study (Discovery Cohort), obtaining tumor and non-tumor adjacent tissue from each of them. Clinical characterization of these patients has been recently reported in detail [7]. The use of human samples for this study was approved by the Ethics Committee of the Reina Sofia University Hospital of Córdoba (Spain) and the study has been conducted following Declaration of Helsinki principles.

Data from a separate cohort of 177 PDAC samples, obtained from the PanCancer study, were also used as Validation Cohort [33]. Clinical and gene expression data from these patients were downloaded from cBioportal [34, 35].

### Cell culture

Two different model cell lines of PDAC were used in functional assays for this study: Capan-2 and MIAPaCa-2 (ATCC, Barcelona, Spain). Capan-2 cells were cultured in McCoy’s 5A Medium (Gibco, Madrid, Spain) supplemented with 10 % fetal bovine serum, 2 mM L-glutamine and 0.2 % of antibiotic/antimycotic. MIAPaCa-2 cells were cultured in Dulbecco’s Modified Eagle’s Medium 4.5 g/L glucose supplemented with 10 % fetal bovine serum, 2.5 % horse serum, 2 mM L-glutamine and 0.2 % of antibiotic/antimycotic. Both cell lines were cultured in a constant humidity 37 °C and 5 % CO_2_ atmosphere. Mycoplasma presence was checked weekly by PCR, as reported in [36].

### RNA isolation, reverse transcription, quantitative PCR and qPCR microfluidic array

Total RNA was isolated from FFPE samples (Discovery Cohort) using Maxwell 16 LEV RNA FFPE Kit (Promega, Madrid, Spain), following manufacturer’s instructions. RNA from cell lines was isolated using TRIzol reagent protocol. RNA was DNAse treated with RNAse-free DNAse kit (Qiagen, Milan, Italy) and it was further quantified using Nanodrop One spectrophotometer (ThermoFisher Scientific, Madrid, Spain).

RNA was reverse transcribed using cDNA First Strand Synthesis kit (ThermoFisher Scientific) with random hexamer primers. Gene expression was quantified using the previously published qPCR protocol [37] based on Brilliant III SYBR Green-QPCR MasterMix (Stratagene, La Jolla, CA, USA) in the Stratagene Mx3000p system.

To simultaneously measure the expression of *EIF4A3* in the Discovery Cohort samples, a dynamic microfluidic qPCR array was used, as previously published by our group [7, 38]. Biomark System and FluidigmVR Real-Time PCR Analysis Software v.3.0.2 and Data Collection Software v.3.1.2 (Fluidigm, South San Francisco, CA, USA) were used to extract and analyze the expression data, using *ACTB, GAPDH* and *HPRT* as housekeeping genes.

### RNA-seq analysis from PDAC samples cohort

An additional Exploration Cohort of 94 samples of tumor tissue from patients with PDAC, comprising clinical and RNA-seq data (described in [7]), was used to explore the associations of *EIF4A3* expression with that of other genes and with the features and patterns of alternative splicing. To this aim, RNA-seq reads were pseudo-aligned and quantified using Salmon and GENCODE v34 version of the human transcriptome. Salmon quantification files were imported to R using Tximeta package. Gene expression was normalized using variance stabilizing transformation (VST) with DESeq2. The patients were classified according to *EIF4A3* expression into three groups using mclust E model by mclust R package.

Alternative splicing events were quantified using SUPPA2 from Salmon quantification files. Briefly, Percent Spliced In (PSI) indexes were quantified for each of the events in the transcriptome in every sample. Delta PSI (ΔPSI) and *p* value were calculated for each of the events comparing high and low *EIF4A3* expression groups. Those events with *p* value < 0.05 were considered as statistically significant for differential alternative splicing analysis.

### Gene silencing using specific siRNA

*EIF4A3* expression was silenced in the cell lines using specific predesigned small interference RNAs (siRNA; #138378, ThermoFisher Scientific). A Negative Control siRNA was also used (#4390843, ThermoFisher Scientific). Specifically, cell lines were transfected using a mix of 30 nM concentration of siRNA and RNAiMax lipofectamine (ThermoFisher Scientific), following manufacturer’s instructions. Cells were detached after 48 h for further assays.

### Proliferation assay

To study proliferation rate in response to *EIF4A3* silencing, 3,000 transfected cells/well were seeded and 10 % resazurin medium was added and incubated for 3 h. Fluorescence (540 nm excitation/ 590 nm emission) was measured at 0, 24, 48 and 72 h at a FlexStation III system (Molecular Devices, Sunnyvale, CA, USA).

### Migration assay

Migration capacity in response to *EIF4A3* silencing was tested using wound healing assay. Briefly, cells were seeded and grown until total confluence and then a wound was made in the well with a pipette tip. Pictures were obtained at 0 and 24 h after wound was done to calculate the area recovered by the cells’ migration. Pictures were analyzed with ImageJ software v.1.51 [39].

### Colonies formation assay

Clonogenic assay was performed to evaluate the cells’ ability to form colonies. Five thousand transfected cells per well were seeded in 6-well plates and were grown for 10 days. Cells were then fixed and stained with Crystal Violet (0.5%) and 6 % formaldehyde solution. Pictures were obtained from the plates, and they were analyzed using ImageJ software v.1.51 [39].

### Tumorspheres formation assay

Tumorspheres formation assay was performed by seeding transfected cells in Corning Costar ultra-low attachment plates (#CLS3473, Merck, Madrid, Spain) in DMEM F-12 medium (#11320033, Gibco) supplemented with EGF (20 ng/ml) (#SRP3027, Sigma-Aldrich, Madrid, Spain) for 10 days. Pictures were obtained from the wells after 10 days of culture and analyzed using ImageJ software v.1.51 [39].

### Xenografted mouse model

A basement membrane extract (#3432-010-01, Trevigen, Gaithersburg, MD, USA) suspension with 1 million MIAPaCa-2 cells was injected subcutaneously in each flank of a 7-weeks-old male athymic BALB/cAnNRj-Foxn1nu mouse (Janvier Labs, Le Genest-Saint-Isle, France). Tumor growth was measured with a digital caliper twice a week. After 15 days, *EIF4A3* and negative control siRNAs were injected in each of the flank tumors of the mice using AteloGene Reagent (#KKN1394, KOKEN Co, Tokyo, Japan). Tumor growth was measured twice a week for two weeks from the injection, when mice were sacrificed. These experiments were performed according to the European-Regulations for Animal-Care under the approval of the University of Cordoba research ethics committees.

### RNA sequencing of silenced cell line

High quality RNA (RNA Integrity Number checked with Agilent BioAnalyzer) from MIAPaCa-2 cells was sequenced after transfecting with *EIF4A3* and negative control siRNAs (n = 3). RNA-seq was performed at the Centre for Genomic Regulation (CRG, Barcelona, Spain). Briefly, ribosomal RNA was depleted, the remaining RNA was fragmented, and library was prepared using Illumina kits. Paired-end sequencing was performed on a HiSeq2500 Illumina instrument to a yield of > 50 million reads per sample. As previously described, FASTQ files were pseudoaligned and quantified using Salmon and gene expression analysis were performed using Tximeta and DESeq2, while alternative splicing analysis were done using SUPPA2.

### Biocomputational and statistical analyses

All statistical analyses from cell lines experiments were performed using Prism 9 software (GraphPad Software, La Jolla, CA, USA). Normality distribution of continuous variables was checked using Shapiro-Wilk test. Mean values were compared using *t*-test, Mann-Whitney *U*-tests or one way ANOVA, depending on the result of data normality check and the number of groups to be compared. Correlation analyses were performed using Pearson tests. Gene expression analyses were performed with R version 4.1.0, using different packages for each analysis: survival_3.2-13 and survminer_0.4.9 for survival analysis, dnet_1.1.7 for gene enrichment analysis, pheatmap_1.0.12 for heatmap representation and ggplot_3.3.5 for graph visualization. Gene Set Enrichment Analyses were performed using GSEA_4.2.3 software. Significance was established at *p* < 0.05.

## RESULTS

### *EIF4A3* is overexpressed in Pancreatic Ductal Adenocarcinoma and associated with poor prognosis

*EIF4A3* expression was measured by RT-qPCR microfluidic array in tumor and non-tumor adjacent tissues from 75 PDAC patients of the Discovery Cohort, showing an overexpression of this gene in tumor tissue at mRNA level (**Fig 1a**). Receiver operating characteristic (ROC) curve analysis confirmed the capacity of *EIF4A3* expression levels to distinguish between tumor and non-tumor tissues, with an area under the curve (AUC) of 0.6129 (**Supp Fig 1**). Interestingly, comparing *EIF4A3* expression in tumor samples from the PanCancer study (TCGA) and healthy pancreas tissue from The Genotype-Tissue Expression (GTEx) confirmed a higher expression of *EIF4A3* in PDAC than in healthy pancreas tissue (**Fig 1b**). Further, increased *EIF4A3* expression was associated to relevant clinical features, such as tumor staging (higher in T4 and in stage III) and metastatic disease (**Fig 1c**). Most importantly, in the PanCancer dataset, *EIF4A3* expression was clearly associated to poorer Overall, Progression Free and Disease Specific survival of patients (**Fig 1d**).

**Fig 1.**
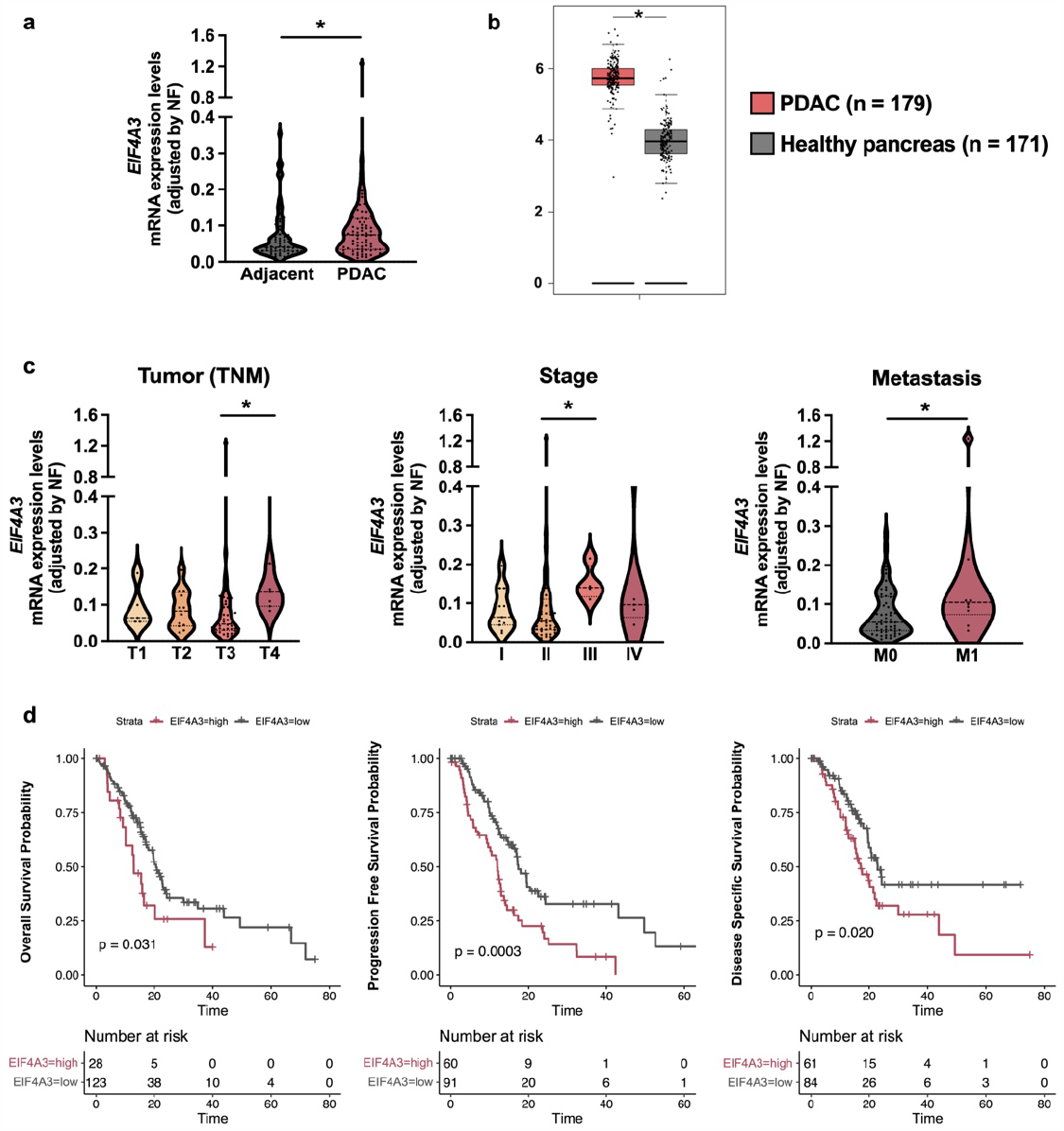
Overexpression of *EIF4A3* in PDAC is associated with malignancy features. **A** *EIF4A3* expression in PDAC tumor tissue compared to non-tumor adjacent tissue as measured by qPCR microfluidic array in the Discovery Cohort. **B** *EIF4A3* expression in The Cancer Genome Atlas (TCGA) PDAC tumors compared to Normal Pancreas expression from The Genotype-Tissue Expression (GTEx). Graphic was obtained by GEPIA web server [57]. **C** Association of *EIF4A3* expression levels in tumor tissue to TNM stage, tumor stage and metastasis in the Discovery Cohort. **D** Association of *EIF4A3* expression levels in tumor tissue to poorer overall, progression free and disease specific survival probability in PDAC patients from the PanCancer study. Data represent mean ± SEM. Asterisks indicate significant differences (*p < 0.05).

### *EIF4A3* expression in PDAC is associated to distinct molecular pathways

To explore the molecular mechanisms associated to *EIF4A3* expression in PDAC, we explored RNA-seq data from another set of 94 tumor samples (Exploration Cohort). GSEA using Cancer Hallmarks Gene set (**Fig 2a**) demonstrated that *EIF4A3* was associated to cell stress responses (UV response, Unfolded Protein response, Apoptosis), metabolic pathways (Glycolysis) and other relevant signaling pathways (MTORC1, TNF-α or MYC). In line with this, enrichment analysis showed a direct correlation of *EIF4A3* mRNA levels in the tumors and the expression of many glycolysis related genes (**Fig 2b**). Furthermore, KEGG pathway analysis (**Fig 2c**) exposed an association between *EIF4A3* expression and a number of metabolic pathways previously related to PDAC [40], from pyridine and purine metabolism to pentose phosphate pathway and to the metabolism of arginine, proline and other amino acids, as well as cell cycle. Interestingly, this analysis also revealed an association of *EIF4A3* expression with that of the spliceosome (**Fig 2c**), and a particularly tight association with that of the other core components of the EJC, *MAGOH, MAGOHB* and *RBM8A* (**Fig 2d**).

**Fig 2.**
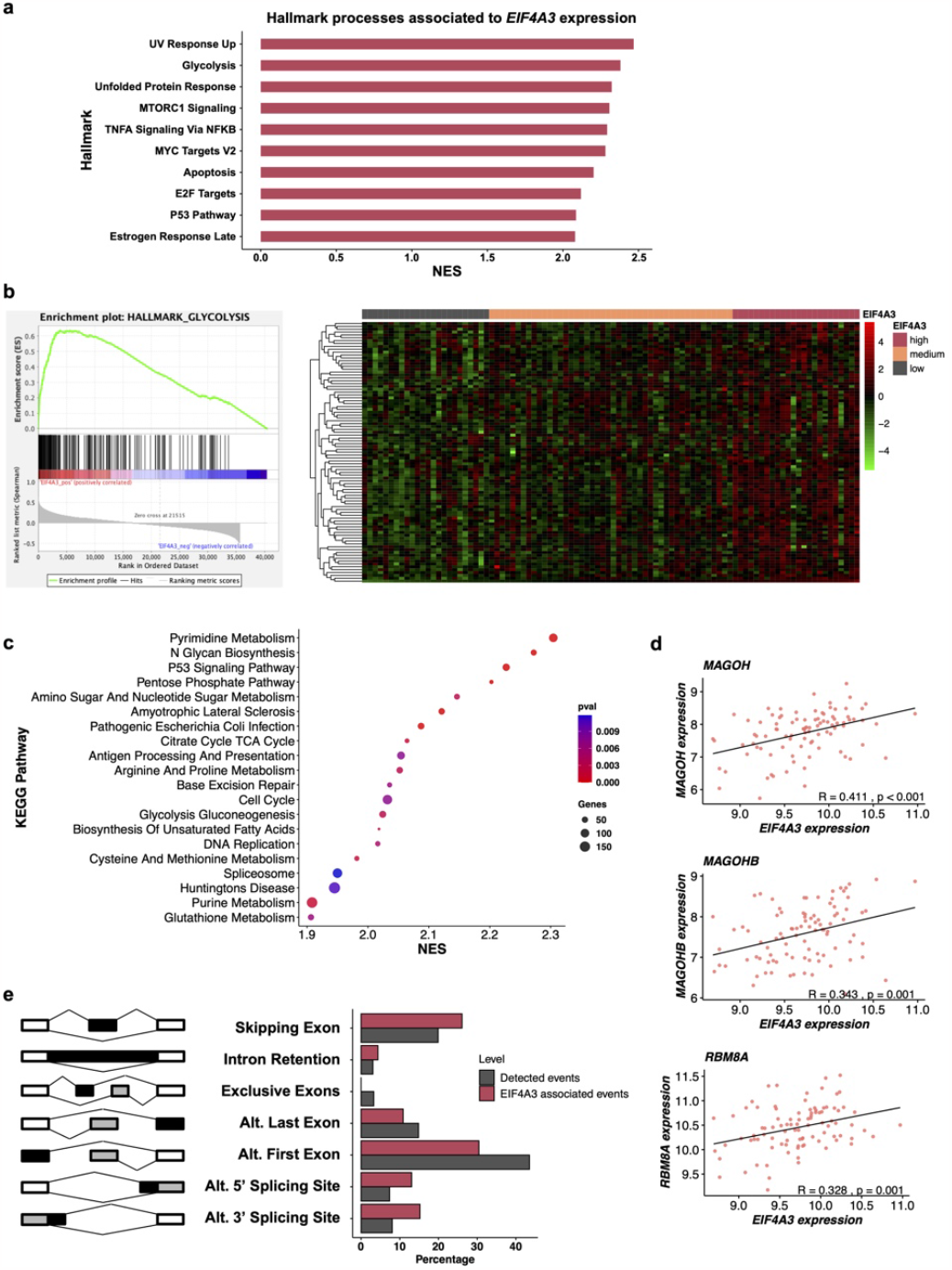
Analysis of *EIF4A3* expression in relation to PDAC molecular landscape in the Exploration Cohort. **A** Analysis of cancer hallmarks significantly associated to *EIF4A3* expression. Normalized enrichment score (NES) is plotted against hallmarks terms. **B** Enrichment plot showing glycolysis hallmark associated to *EIF4A3* expression (left) and non-hierarchical heatmap showing the expression of glycolysis genes in low, intermediate, and high *EIF4A3* expression PDAC samples (right). **C** Analysis of KEGG pathways significantly associated to *EIF4A3* expression. NES is plotted against KEGG pathways, dot size represents the genes hits and dot colors represents *p* value of that pathway. **D** *EIF4A3* expression Pearson correlation to the other components of the Exon Junction Complex. **E** Bar plot showing the frequency from each alternative splicing event type detected in PDAC tumors (grey) and from those statistically associated to *EIF4A3* expression (red).

Based on these correlations with the expression of multiple spliceosome components, we posited that the levels of *EIF4A3* expression could also be associated to distinctive patterns of alternative splicing. Interestingly, when comparing differential alternative splicing events between tumors expressing high or low *EIF4A3* levels, we found an overrepresentation of skipping exons and alternative 5’ and 3’ splicing sites events, whereas alternative first and last exons events were less frequent (**Fig 2e**). These results suggest a link between alteration of *EIF4A3* expression and alternative splicing dysregulation in PDAC.

### *EIF4A3* silencing decreases malignant functional features *in vitro* and *in vivo*

Based on the results described above, we hypothesized that decreasing *EIF4A3* expression could exert antitumor effects on PDAC. To test this notion, we used two model cell lines, Capan-2 and MIAPaCa-2, where *EIF4A3* was silenced using either a specific siRNA (**Supp Fig 2**) or a negative control siRNA (scramble). As illustrated in **Fig 3a**, *EIF4A3* silencing decreased proliferation in both cell lines, although it was noteworthy that the effect was more apparent and time-dependent in the more aggressive and poorly differentiated MIAPaCa-2 cells, than in the well differentiated Capan-2 cells, which only reduced their proliferation at 72 h. Migration capacity of MIAPaCa-2 cells was also assessed in response to *EIF4A3* silencing by wound healing assay, where a clear decrease in the recovery of the wound at 24 h was observed (**Fig 3b**). Capan-2 cells could not be tested by this method because of their growing pattern at full confluence, which does not allow to use this assay.

**Fig 3.**
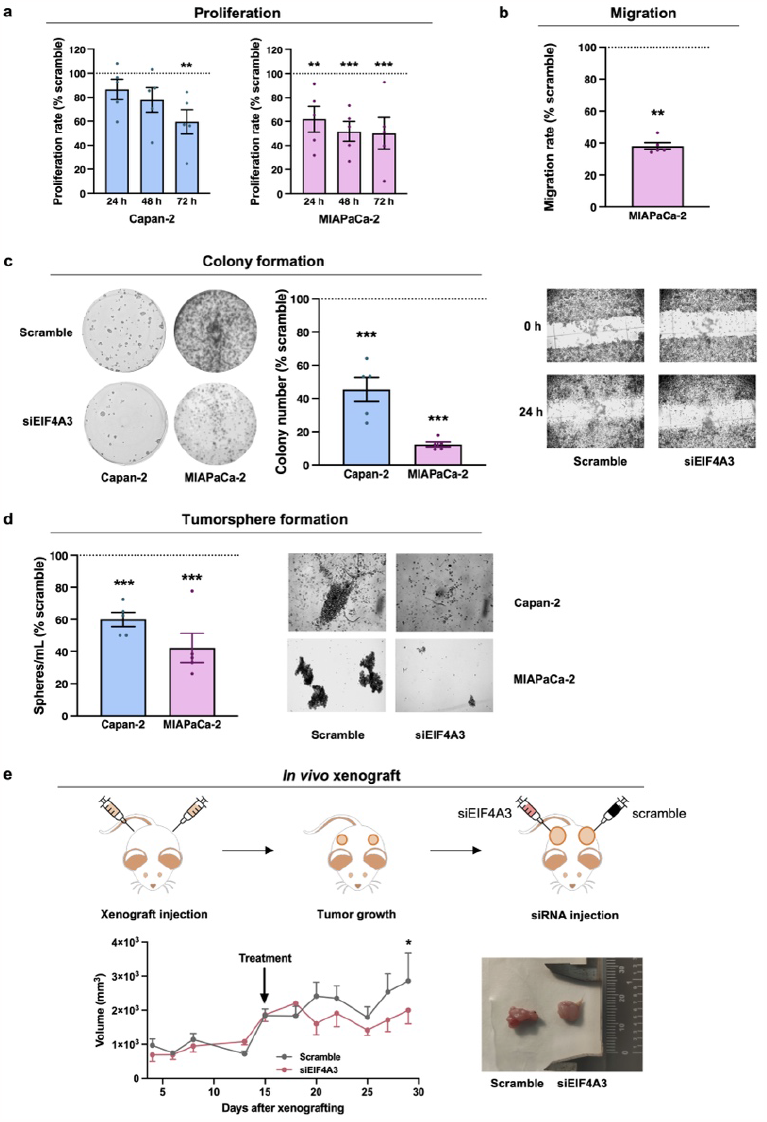
Functional effects of EIF4A3 targeting in PDAC models. **A** Proliferation rates of Capan-2 and MIAPaCa-2 cell lines in response to *EIF4A3* silencing at 24, 48 and 72 h compared to scramble silenced cells (n = 5). **B** Migration rate of MIAPaCa-2 cells in response to *EIF4A3* silencing compared to scramble silenced cells set at 100% (n = 5). **C** Colony formation capacity of Capan-2 and MIAPaCa-2 cells in response to *EIF4A3* silencing compared to scramble silenced cells set at 100% (n = 5). **D** Tumorsphere formation capacity of Capan-2 and MIAPaCa-2 cells in response to *EIF4A3* silencing compared to scramble silenced cells set at 100% (n = 5). **E** EIF4A3 targeting reduces tumor volume *in vivo* (n = 6). At top panel, scheme of the xenograft experiment carried out; at bottom-left panel, volume of the tumors (*EIF4A3* and scramble silenced groups) during the full experiment; at bottom-right panel, representative picture of the tumors after the mice sacrifice. Data represents mean ± SEM. Asterisks indicate significant differences (\**p* < 0.05; \*\**p* < 0.01; \*\*\**p* < 0.001).

Colony formation was also assessed after *EIF4A3* silencing (**Fig 3c**), and a profound decrease was observed in both cell lines compared to scramble controls. Once again, the effect was stronger in MIAPaCa-2 cells, suggesting a particular link between this gene and stem cell properties in highly aggressive cells. Similar to colony formation, when tumorspheres formation capacity was tested in both cell lines, a decrease in the density of spheres formation was observed in response to *EIF4A3* silencing, when compared to scramble controls (**Fig 3d**).

To further evaluate if the effects of *EIF4A3* silencing were also recapitulated *in vivo*, a xenograft model was generated by subcutaneous injection of MIAPaCa-2 cells in both flanks of athymic mice (n = 6). In each of the two tumors formed, we injected *EIF4A3* or scramble siRNAs as a negative control. After siRNAs injection, a decrease in tumor volume was observed only in tumors that received the *EIF4A3* silencing treatment (**Fig 3e**). The difference in tumor volume increased over time and became statistically significant at 14 days. These results indicate that *EIF4A3* silencing has antitumor capacity not only *in vitro* but also *in vivo*.

### *EIF4A3* silencing alters gene expression of key molecular pathways

To explore the mechanistic underpinnings of *EIF4A3* silencing, we carried out an RNA-seq experiment with MIAPaCa-2 cells transfected with either *EIF4A3* or scramble siRNAs (n = 3). A robust (> 50%) *EIF4A3* silencing was confirmed after the sequencing (**Fig 4a**). Overall, 191 genes were upregulated, and 138 genes were downregulated in *EIF4A3* silenced cells (**Fig 4b**). To identify the molecular pathways these genes belong to, we performed two analyses. First, Hallmarks Gene Set Analysis (**Fig 4c**) showed that there was an upregulation of MYC targets and oxidative phosphorylation genes, whereas other pathways like pancreas beta cells genes, the relevant KRAS signaling and TNF-α signaling sets, or immunity related gene sets like allograft rejection and complement, were downregulated. Further, KEGG pathways analysis revealed that after *EIF4A3* silencing, the main altered pathways were related to RNA processing and metabolic pathways (**Fig 4d**). Overall, the main pathways associated to *EIF4A3* expression observed previously in the analysis of human samples (**Fig 2**) and those altered after its silencing in MIAPaCa-2 cells substantially overlapped, as the latter related to carbon metabolism (oxidative phosphorylation) (**Fig 4e**) and spliceosome (including EJC components *MAGOH* and *MAGOHB*) (**Fig 4f**), which were both upregulated after *EIF4A3* silencing; and TNF-alpha signaling (**Fig 4g**), which was repressed after *EIF4A3* silencing.

**Fig 4.**
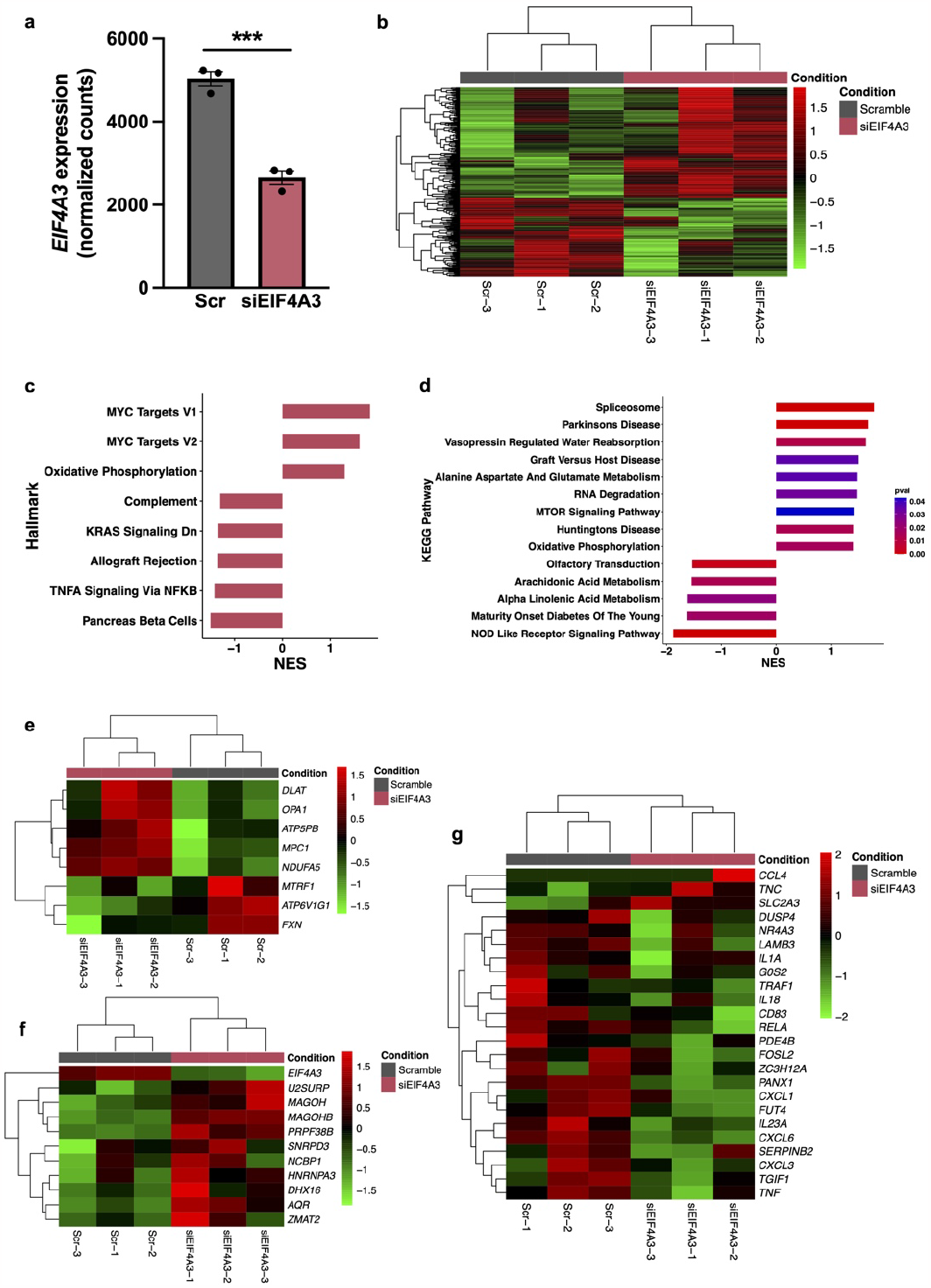
Changes in the transcriptional profile of MIAPaCa-2 cells in response to *EIF4A3* silencing. **A** Normalized *EIF4A3* expression in scramble vs *EIF4A3* silenced MIAPaCa-2 cells (n = 3). **B** Heatmap showing the differentially expressed genes (DEG) by DESeq2 in scramble vs *EIF4A3* silenced MIAPaCa-2 cells. **C** Analysis of Hallmarks altered due to *EIF4A3* silencing in MIAPaCa-2 cells. NES is positive for hallmarks enriched in *EIF4A3* silenced cells and NES is negative for those enriched in scramble silenced cells. **D** Analysis of KEGG pathways altered due to *EIF4A3* silencing in MIAPaCa-2 cells. NES is positive for hallmarks enriched in *EIF4A3* silenced cells and NES is negative for those enriched in scramble silenced cells. Bars color represents *p* value for each pathway. **E-G** Heatmap showing the DEG expression from the oxidative phosphorylation (**E**), the spliceosome (**F**) and the TNF-α (**G**) pathways.

### *EIF4A3* silencing modifies alternative splicing pattern in PDAC cells

Analysis of alternative splicing in RNA-seq data from MIAPaCa-2 transfected cells revealed that 280 genes were differentially spliced when *EIF4A3* was silenced, while, in contrast, most of them did not change in their expression levels (**Fig 5a**). Remarkably, there were 535 alternative splicing events that changed their PSI when *EIF4A3* was silenced (**Fig 5b**). Those events mainly belong to skipping exon and alternative first exon types, according to SUPPA2 analysis. Also, the pattern of splicing events that changed after *EIF4A3* silencing (*p* < 0.05) was different from the other alternative splicing events detected, being skipping exon or alternative 3’ splice site overrepresented among them, whereas alternative first and last exons were less frequent (**Fig 5c**). Additional enrichment analyses were performed to identify which genes were affected by the changes in alternative splicing pattern. Biological process Gene Ontology (GO) analysis showed that silencing *EIF4A3* mainly affected RNA translation and apoptosis related genes (**Fig 5d**). Collectively, these results strongly suggest a relevant role for EIF4A3 in the selective regulation of alternative splicing events influencing capital biological functions in PDAC.

**Fig 5.**
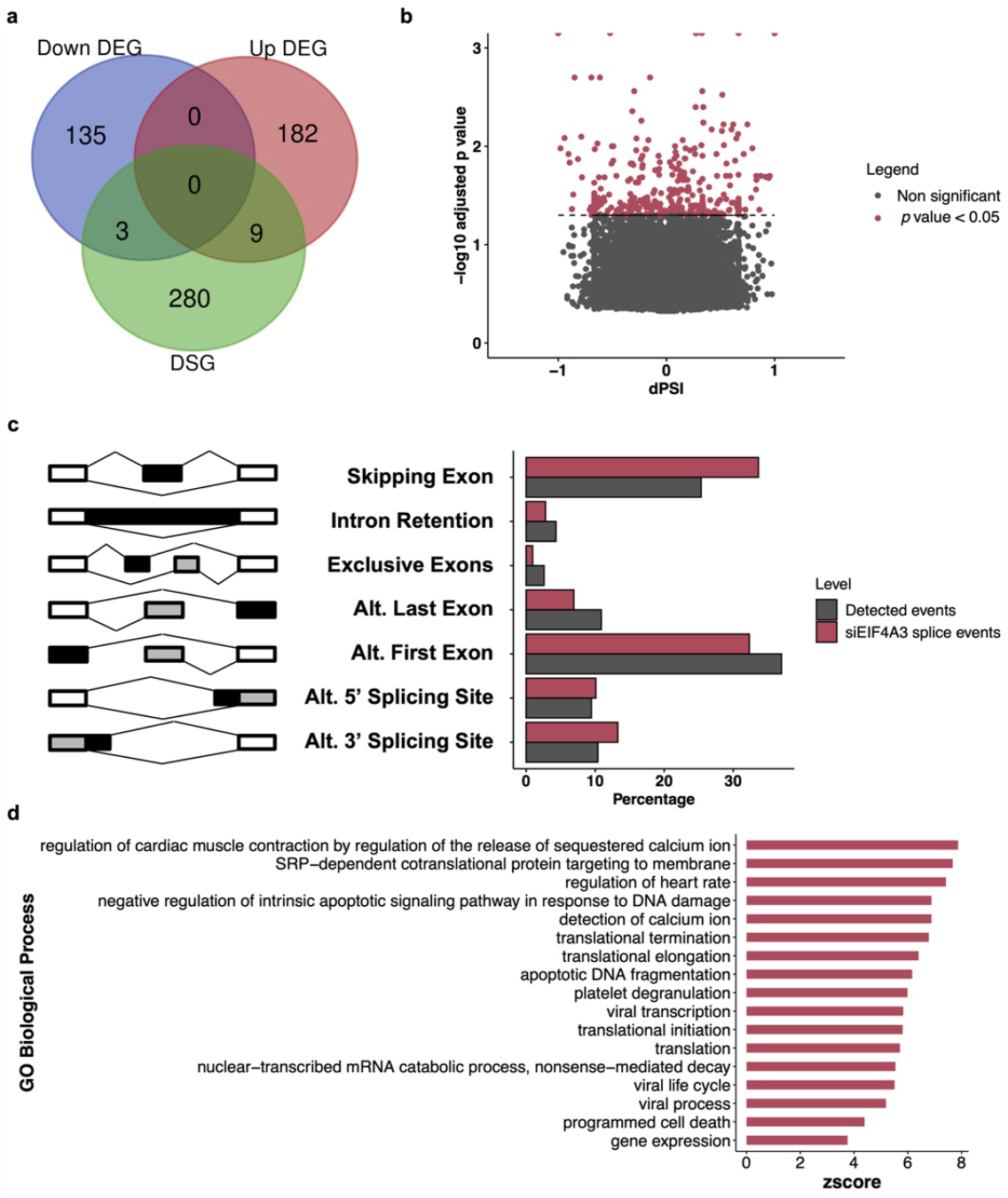
Changes in alternative splicing in MIAPaCa-2 cells associated to *EIF4A3* silencing. **A** Venn diagram showing the intersection among the downregulated DEG, upregulated DEG and the differentially spliced genes (DSG) after *EIF4A3* silencing. **B** Volcano plot showing the differentially alternative splice events after *EIF4A3* silencing in MIAPaCa-2 cells. Differential Percent Splice In (dPSI) from each event is plotted against the logarithm of its *p* value. Those events with *p* < 0.05 are colored. **C** Bar plot showing the frequency from each alternative splicing event type detected in MIAPaCa-2 cells (grey) and from those statistically differential after *EIF4A3* silencing (red). **D** Analysis of Biological Process Gene Ontology (GO-BP) pathways enriched for the genes that were alternatively spliced after *EIF4A3* silencing (*p* < 0.05). Zscore is plotted for each of the GO-BP pathways. Only those pathways with adjusted *p* value < 0.01 are represented.

## DISCUSSION

Pioneer reports showing two decades ago an altered expression of abnormal CD44 variants and other “spliceoforms” in PDAC [41] heralded the current bloom of experimental evidence supporting a major role of splicing dysregulation in this deadly cancer [12]. Subsequent work reinforced this notion by showing that the splicing machinery itself is dysregulated in PDAC, and can thereby contribute to trigger some of its typical pathological features, from its precursor pancreatitis [11], to increased cell proliferation and metastasis [7, 10], reviewed in [12]. Further, recent evidence is helping to dissect key molecular players that interact with and mediate the pathological impact of dysregulated splicing components, such as hnRNPK [9], SF3B1 [7], RBFOX2 [10], SRSF1 [11], and others, including core oncogenic hallmarks of PDAC like KRAS and TP53 [2]. In this context, there is still a notable dearth of knowledge on the potential role in PDAC of molecular players and mechanisms that, like the EJC component EIF4A3, are essential to maintain RNA homeostasis and splicing itself, and are involved in cancer [17–19]. In this scenario, our present study provides original evidence that EIF4A3 is altered in PDAC, where it associates to clinical and molecular features, and may serve a pathological role, mechanistically linked to the alteration of multiple pathways related to both metabolic routes and splicing and RNA biology.

Our initial interest on EIF4A3 was sparked by its involvement in splicing and by the growing evidence of its role in several cancers, which contrasted with the limited information available in PDAC. In fact, detailed studies have shown a relevant role of two members of the EIF4A family, EIF4A1 and EIF4A2, in PDAC, where they seem to participate in the translational control of pancreatic tumor metabolism, a dependency potentially exploitable therapeutically [24, 25]. A similar metabolic link was reported for EIF4A1 and a related family member, EIF4E, in relation to phosphoglycerate dehydrogenase and translation initiation in PDAC [42]. However, few studies have explored EIF4A3 in PDAC and were focused on its interaction with other molecular players [27, 28]. In contrast, a growing number of studies are providing detailed experimental support for a relevant role of EIF4A3 in various cancers, including hepatocellular carcinoma [30], breast cancer [31] cervical cancer [32] or glioblastoma [43, 44] (reviewed in [18]). Although the underlying mechanisms proposed to mediate EIF4A3 differ for each tumor type investigated, there is a common trend involving RNA-related molecules and processes [18]. In PDAC, EIF4A3 has been proposed to participate in the biogenesis of an oncogenic circRNA (circRNF13) in relation to the hypoxia-inducible factor-1 (HIF-1), a key player in aerobic glycolysis [28]. In turn, circRNF13 would promote cell proliferation, angiogenesis, invasion and glycolysis in PDAC cells and mice models by acting as a sponge for miR-654-3p and thereby augmenting the levels of pyruvate dehydrogenase kinase 3 (PDK3) [28]. On a distinct but related setting, EIF4A3 was recently proposed to mediate the oncogenic effect on PDAC of the long non-coding RNA LINC01232, which, after being induced by the ubiquitous transcription factor SP1, would recruit EIF4A3 to upregulate expression of the transmembrane 9 superfamily member 2 (TM9SF2) by enhancing its mRNA stability [27]. These studies document EIF4A3 as a relevant conduit mediating the actions of RNA-related actors in PDAC. However, they could not provide a more general vision of its role in this cancer.

By specifically focusing on the role and implications of EIF4A3 itself, our present work uncovered original evidence suggestive of an overarching regulatory role of this factor in PDAC, providing novel insights on its plausible influence at crossroads of metabolic routes, splicing and RNA biology. Indeed, *EIF4A3* expression levels showed tight links to glycolysis and several of its associated metabolic pathways, which are well known pivotal actors for PDAC oncogenesis and progression [2], not only in association studies in human PDAC samples but also after experimentally-driven causative modifications in cell lines. These findings are in line with reports on the ability of EIF4A3 to enhance biogenesis of glycolysis-inducing circRNAs in PDAC [28] and prostate cancer [45]. Interestingly, glycolysis activation in PDAC is known to be related to c-MYC [46], which has been proven to be persistently and aberrantly activated in this cancer [47, 48]. In turn, MYC hyperactivation is associated to dysregulation of key components of the splicing machinery in PDAC like SRSF1 [11]. In this scenario, our data showing a close correlation of *EIF4A3* expression with that of MYC- and glycolysis-related pathways in PDAC reinforce the emerging connection of MYC hyperactivation with splicing and metabolic dysregulation in cancer [11, 45, 49]. In a related metabolic context, we found that *EIF4A3* silencing enhanced expression of oxidative phosphorylation genes, suggesting a change in central carbon metabolism, which is known to be altered in PDAC, where it has been recently proposed as an anti-mitochondrial respiration therapy in specific patients [50]. Of note, our data also unveils that *EIF4A3* expression in PDAC is closely associated to several pathways classically linked to stress response, from apoptosis to UV response or Unfolded Protein response, which nicely fits a general (not just cancer-related) stress-linked function recently proposed for this molecule [51]. Nonetheless, the associations of EIF4A3 expression levels were not circumscribed to metabolic or splicing pathways, but also involved other pivotal factors in PDAC, such as TNF-α, which has been proposed as a potential therapy in this pathology [52, 53]. Specifically, *EIF4A3* silencing decreased the expression of TNF-α signaling pathway genes, including TNF-α itself, thereby implying putative interactions with PDAC microenvironment and metastasis [52, 54].

A particular novelty of our study is the discovery of the clear influence of *EIF4A3* expression on the global patterns of splicing in PDAC and how this translates to selective changes in specific gene families. Actually, the lack of information in this regard in PDAC was somewhat surprising, given the known roles of EIF4A3, as EJC component, in linking splicing with other processes like RNA export, NMD, and global RNA homeostasis [18, 55]. Interestingly, our data revealed that the increased expression levels of *EIF4A3* in human PDAC samples is associated to markedly altered patterns of alternative splicing. Moreover, experimental *EIF4A3* silencing in PDAC cells, rather than altering gene expression patterns, caused profound changes in alternative splicing profiles, which, in keeping with previous reports on EIF4A3 and the EJC, particularly involved skipping exons events [56]. Detailed analysis of the molecular pathways affected by alternative splicing changes mediated by *EIF4A3* downregulation suggested a putative involvement of this factor in the regulation of cellular processes essential in cancer progression, such as those affecting apoptosis, NMD and RNA translation [17], [22]. These observations suggest that EIF4A3 would not only participate in the regulation of correct RNA metabolism but may also influence PDAC cell survival, disclosing its potential as an actionable target. In line with this notion, our *in vitro* and *in vivo* results demonstrated the functional therapeutic benefits of targeting *EIF4A3*. Specifically, its silencing *in vitro* reduced aggressiveness parameters (proliferation and migration rates) of PDAC cell lines, and reduced stemness functional parameters (colony and tumorspheres formation). Likewise, *EIF4A3* silencing blunted tumor growth in a preclinical model (MIAPaCa-2 derived xenografted mouse). Our results are in close agreement with recent reports showing that targeting EIF4A3 can reduce tumor growth or aggressiveness features in other cancer types [30–32, 43–45], reviewed in [18], thereby leveraging its potential as an actionable target in cancer.

In conclusion, our results indicate that the EJC component *EIF4A3* is abnormally overexpressed in PDAC, where it associates to malignant clinical features and poor patient survival. In line with the increasingly ample and relevant functions reported for EIF4A3, we found that this factor may play multiple roles in PDAC, involving from key metabolic pathways (glycolysis, oxidative phosphorylation) to RNA translation, NMD and, particularly splicing, thereby suggesting its potential as an overarching hub for the homeostasis of RNA biology. Finally, experimental *in vitro* and *in vivo* targeting of EIF4A3 decreased aggressiveness features and tumor growth of PDAC cells, providing primary evidence of its potential as a candidate therapeutic target in this dismal cancer.

## STATEMENTS AND DECLARATIONS

### Authors contributions

**RBE**: Investigation, Formal analysis, Software, Data Curation, Writing - Original Draft; **EAP**: Investigation, Formal analysis, Data Curation, Writing – Review & Editing; **MTMM**: Investigation, Writing – Review & Editing; **VGV**: Investigation, Writing – Review & Editing; **MESF**: Resources, Writing – Review & Editing; **AM**: Investigation, Resources, Writing – Review & Editing; **JLLC**: Investigation, Writing – Review & Editing; **CB**: Investigation, Writing – Review & Editing; **MDG**: Investigation, Writing – Review & Editing; **RTL**: Investigation, Resources, Writing – Review & Editing; **RML**: Investigation, Writing – Review & Editing; **AS**: Investigation, Resources, Writing – Review & Editing; **ÁAS**: Resources, Writing – Review & Editing; **SPA**: Investigation, Writing – Review & Editing; **AIC**: Investigation, Supervision, Writing – Review & Editing; **JPC**: Conceptualization, Investigation, Supervision, Resources, Writing – Original Draft.

### Data availability

The datasets used and/or analyzed during the current study are available from the corresponding author upon reasonable request.

### Ethics approval

This study was approved by the Ethics Committee of the Reina Sofia University Hospital and the Declaration of Helsinki guidelines were followed. Mice xenograft experiments were performed according to the European-Regulations for Animal-Care under the approval of the University of Cordoba research ethics committees.

### Consent to participate

Informed consent documentation was obtained from each of the patients involved in the study.

## Acknowledgements

We thank all patients and their families for generously donating the samples studies in this work.

## SUPPLEMENTARY FIGURES LEGENDS

**Supp Fig 1** Receiver operating characteristic (ROC) curve analysis of *EIF4A3* expression in tumor and non-tumor adjacent tissue distinction in the Discovery Cohort.

**Supp Fig 2** *EIF4A3* silencing validation in Capan-2 and MIAPaCa-2 cell lines measured by qPCR. Expression is represented as percentage compared to the Scramble silenced cells, and it was normalized using *ACTB* as housekeeping gene. Data represents mean ± SEM. Asterisks indicate significant differences (\**p* < 0.05).

## REFERENCES

1. Grossberg AJ, Chu LC, Deig CR, et al (2020) Multidisciplinary standards of care and recent progress in pancreatic ductal adenocarcinoma. CA Cancer J Clin 70:375–403. 10.3322/caac.21626

2. Halbrook CJ, Lyssiotis CA, Pasca di Magliano M, Maitra A (2023) Pancreatic cancer: Advances and challenges. Cell 186:1729–1754. 10.1016/j.cell.2023.02.014

3. Sung H, Ferlay J, Siegel RL, Laversanne M, Soerjomataram I, Jemal A, Bray F (2021) Global Cancer Statistics 2020: GLOBOCAN Estimates of Incidence and Mortality Worldwide for 36 Cancers in 185 Countries. CA Cancer J Clin 71:209–249. 10.3322/caac.21660

4. Bailey P, Chang DK, Nones K, et al (2016) Genomic analyses identify molecular subtypes of pancreatic cancer. Nature 531:47–52. 10.1038/nature16965

5. Waddell N, Pajic M, Patch A-M, et al (2015) Whole genomes redefine the mutational landscape of pancreatic cancer. Nature 518:495–501. 10.1038/nature14169

6. Connor AA, Gallinger S (2022) Pancreatic cancer evolution and heterogeneity: integrating omics and clinical data. Nat Rev Cancer 22:131–142. 10.1038/s41568-021-00418-1

7. Alors-Perez E, Blázquez-Encinas R, Alcalá S, et al (2021) Dysregulated splicing factor SF3B1 unveils a dual therapeutic vulnerability to target pancreatic cancer cells and cancer stem cells with an anti-splicing drug. J Exp Clin Cancer Res CR 40:382. 10.1186/s13046-021-02153-9

8. Wang J, Dumartin L, Mafficini A, et al (2017) Splice variants as novel targets in pancreatic ductal adenocarcinoma. Sci Rep 7:2980. 10.1038/s41598-017-03354-z

9. Escobar-Hoyos LF, Penson A, Kannan R, et al (2020) Altered RNA Splicing by Mutant p53 Activates Oncogenic RAS Signaling in Pancreatic Cancer. Cancer Cell 38:198–211.e8. 10.1016/j.ccell.2020.05.010

10. Jbara A, Lin K-T, Stossel C, et al (2023) RBFOX2 modulates a metastatic signature of alternative splicing in pancreatic cancer. Nature 617:147–153. 10.1038/s41586-023-05820-3

11. Wan L, Lin K-T, Rahman MA, Ishigami Y, Wang Z, Jensen MA, Wilkinson JE, Park Y, Tuveson DA, Krainer AR (2023) Splicing Factor SRSF1 Promotes Pancreatitis and KRASG12D-Mediated Pancreatic Cancer. Cancer Discov 13:1678–1695. 10.1158/2159-8290.CD-22-1013

12. Alors-Pérez E, Pedraza-Arevalo S, Blázquez-Encinas R, Moreno-Montilla MT, García-Vioque V, Berbel I, Luque RM, Sainz B, Ibáñez-Costa A, Castaño JP (2023) Splicing alterations in pancreatic ductal adenocarcinoma: a new molecular landscape with translational potential. J Exp Clin Cancer Res CR 42:282. 10.1186/s13046-023-02858-z

13. Marasco LE, Kornblihtt AR (2022) The physiology of alternative splicing. Nat Rev Mol Cell Biol. 10.1038/s41580-022-00545-z

14. Wright CJ, Smith CWJ, Jiggins CD (2022) Alternative splicing as a source of phenotypic diversity. Nat Rev Genet 23:697–710. 10.1038/s41576-022-00514-4

15. Bradley RK, Anczuków O (2023) RNA splicing dysregulation and the hallmarks of cancer. Nat Rev Cancer 23:135–155. 10.1038/s41568-022-00541-7

16. Wang E, Aifantis I (2020) RNA Splicing and Cancer. Trends Cancer 6:631–644. 10.1016/j.trecan.2020.04.011

17. Wolin SL, Maquat LE (2019) Cellular RNA surveillance in health and disease. Science 366:822–827. 10.1126/science.aax2957

18. Ye J, She X, Liu Z, He Z, Gao X, Lu L, Liang R, Lin Y (2021) Eukaryotic Initiation Factor 4A-3: A Review of Its Physiological Role and Involvement in Oncogenesis. Front Oncol 11:712045. 10.3389/fonc.2021.712045

19. Woodward LA, Mabin JW, Gangras P, Singh G (2017) The exon junction complex: a lifelong guardian of mRNA fate. Wiley Interdiscip Rev RNA 8:. 10.1002/wrna.1411

20. Xue C, Gu X, Li G, Bao Z, Li L (2021) Expression and Functional Roles of Eukaryotic Initiation Factor 4A Family Proteins in Human Cancers. Front Cell Dev Biol 9:711965. 10.3389/fcell.2021.711965

21. Gehring NH, Kunz JB, Neu-Yilik G, Breit S, Viegas MH, Hentze MW, Kulozik AE (2005) Exon-junction complex components specify distinct routes of nonsense-mediated mRNA decay with differential cofactor requirements. Mol Cell 20:65–75. 10.1016/j.molcel.2005.08.012

22. Kanellis DC, Espinoza JA, Zisi A, et al (2021) The exon-junction complex helicase eIF4A3 controls cell fate via coordinated regulation of ribosome biogenesis and translational output. Sci Adv 7:eabf7561. 10.1126/sciadv.abf7561

23. Li Y, Ren S, Xia J, Wei Y, Xi Y (2020) EIF4A3-Induced circ-BNIP3 Aggravated Hypoxia-Induced Injury of H9c2 Cells by Targeting miR-27a-3p/BNIP3. Mol Ther Nucleic Acids 19:533–545. 10.1016/j.omtn.2019.11.017

24. Müller D, Shin S, Goullet de Rugy T, et al (2019) eIF4A inhibition circumvents uncontrolled DNA replication mediated by 4E-BP1 loss in pancreatic cancer. JCI Insight 4:e121951. 10.1172/jci.insight.121951

25. Chan K, Robert F, Oertlin C, et al (2019) eIF4A supports an oncogenic translation program in pancreatic ductal adenocarcinoma. Nat Commun 10:5151. 10.1038/s41467-019-13086-5

26. Singh K, Lin J, Lecomte N, et al (2021) Targeting eIF4A Dependent Translation of KRAS Signaling Molecules. Cancer Res 81:2002–2014. 10.1158/0008-5472.CAN-20-2929

27. Li Q, Lei C, Lu C, Wang J, Gao M, Gao W (2019) LINC01232 exerts oncogenic activities in pancreatic adenocarcinoma via regulation of TM9SF2. Cell Death Dis 10:698. 10.1038/s41419-019-1896-3

28. Zhao Q, Zhu Z, Xiao W, et al (2022) Hypoxia-induced circRNF13 promotes the progression and glycolysis of pancreatic cancer. Exp Mol Med 54:1940–1954. 10.1038/s12276-022-00877-y

29. Xia Q, Kong X-T, Zhang G-A, Hou X-J, Qiang H, Zhong R-Q (2005) Proteomics-based identification of DEAD-box protein 48 as a novel autoantigen, a prospective serum marker for pancreatic cancer. Biochem Biophys Res Commun 330:526–532. 10.1016/j.bbrc.2005.02.181

30. López-Cánovas JL, Hermán-Sánchez N, Moreno-Montilla MT, et al (2022) Spliceosomal profiling identifies EIF4A3 as a novel oncogene in hepatocellular carcinoma acting through the modulation of FGFR4 splicing. Clin Transl Med 12:e1102. 10.1002/ctm2.1102

31. Liu Q, Dong H (2021) EIF4A3-mediated hsa_circ_0088088 promotes the carcinogenesis of breast cancer by sponging miR-135-5p. J Biochem Mol Toxicol 35:e22909. 10.1002/jbt.22909

32. Yang M, Hu H, Wu S, Ding J, Yin B, Huang B, Li F, Guo X, Han L (2022) EIF4A3-regulated circ_0087429 can reverse EMT and inhibit the progression of cervical cancer via miR-5003-3p-dependent upregulation of OGN expression. J Exp Clin Cancer Res CR 41:165. 10.1186/s13046-022-02368-4

33. Cancer Genome Atlas Research Network (2017) Integrated Genomic Characterization of Pancreatic Ductal Adenocarcinoma. Cancer Cell 32:185–203.e13. 10.1016/j.ccell.2017.07.007

34. Cerami E, Gao J, Dogrusoz U, et al (2012) The cBio cancer genomics portal: an open platform for exploring multidimensional cancer genomics data. Cancer Discov 2:401–404. 10.1158/2159-8290.CD-12-0095

35. Gao J, Aksoy BA, Dogrusoz U, et al (2013) Integrative analysis of complex cancer genomics and clinical profiles using the cBioPortal. Sci Signal 6:1. 10.1126/scisignal.2004088

36. Uphoff CC, Drexler HG (2005) Detection of mycoplasma contaminations. Methods Mol Biol Clifton NJ 290:13–23. 10.1385/1-59259-838-2:013

37. Ibáñez-Costa A, López-Sánchez LM, Gahete MD, et al (2017) BIM-23A760 influences key functional endpoints in pituitary adenomas and normal pituitaries: molecular mechanisms underlying the differential response in adenomas. Sci Rep 7:42002. 10.1038/srep42002

38. Pedraza-Arevalo S, Alors-Pérez E, Blázquez-Encinas R, et al (2022) Spliceosomic dysregulation unveils NOVA1 as a candidate actionable therapeutic target in pancreatic neuroendocrine tumors. Transl Res J Lab Clin Med S1931-5244(22)00170–0. 10.1016/j.trsl.2022.07.005

39. Schindelin J, Arganda-Carreras I, Frise E, et al (2012) Fiji: an open-source platform for biological-image analysis. Nat Methods 9:676–682. 10.1038/nmeth.2019

40. Guillaumond F, Leca J, Olivares O, et al (2013) Strengthened glycolysis under hypoxia supports tumor symbiosis and hexosamine biosynthesis in pancreatic adenocarcinoma. Proc Natl Acad Sci U S A 110:3919–3924. 10.1073/pnas.1219555110

41. Seiter S, Arch R, Reber S, Komitowski D, Hofmann M, Ponta H, Herrlich P, Matzku S, Zöller M (1993) Prevention of tumor metastasis formation by anti-variant CD44. J Exp Med 177:443–455. 10.1084/jem.177.2.443

42. Ma X, Li B, Liu J, Fu Y, Luo Y (2019) Phosphoglycerate dehydrogenase promotes pancreatic cancer development by interacting with eIF4A1 and eIF4E. J Exp Clin Cancer Res CR 38:66. 10.1186/s13046-019-1053-y

43. Wang R, Zhang S, Chen X, Li N, Li J, Jia R, Pan Y, Liang H (2018) EIF4A3-induced circular RNA MMP9 (circMMP9) acts as a sponge of miR-124 and promotes glioblastoma multiforme cell tumorigenesis. Mol Cancer 17:166. 10.1186/s12943-018-0911-0

44. Tang W, Wang D, Shao L, et al (2020) LINC00680 and TTN-AS1 Stabilized by EIF4A3 Promoted Malignant Biological Behaviors of Glioblastoma Cells. Mol Ther Nucleic Acids 19:905–921. 10.1016/j.omtn.2019.10.043

45. Jiang X, Guo S, Wang S, Zhang Y, Chen H, Wang Y, Liu R, Niu Y, Xu Y (2022) EIF4A3-Induced circARHGAP29 Promotes Aerobic Glycolysis in Docetaxel-Resistant Prostate Cancer through IGF2BP2/c-Myc/LDHA Signaling. Cancer Res 82:831–845. 10.1158/0008-5472.CAN-21-2988

46. Ala M (2022) Target c-Myc to treat pancreatic cancer. Cancer Biol Ther 23:34–50. 10.1080/15384047.2021.2017223

47. Wirth M, Schneider G (2016) MYC: A Stratification Marker for Pancreatic Cancer Therapy. Trends Cancer 2:1–3. 10.1016/j.trecan.2015.12.002

48. Schleger C, Verbeke C, Hildenbrand R, Zentgraf H, Bleyl U (2002) c-MYC activation in primary and metastatic ductal adenocarcinoma of the pancreas: incidence, mechanisms, and clinical significance. Mod Pathol Off J U S Can Acad Pathol Inc 15:462–469. 10.1038/modpathol.3880547

49. Urbanski L, Brugiolo M, Park S, Angarola BL, Leclair NK, Yurieva M, Palmer P, Sahu SK, Anczuków O (2022) MYC regulates a pan-cancer network of co-expressed oncogenic splicing factors. Cell Rep 41:111704. 10.1016/j.celrep.2022.111704

50. Masoud R, Reyes-Castellanos G, Lac S, et al (2020) Targeting Mitochondrial Complex I Overcomes Chemoresistance in High OXPHOS Pancreatic Cancer. Cell Rep Med 1:100143. 10.1016/j.xcrm.2020.100143

51. Mazloomian A, Araki S, Ohori M, et al (2019) Pharmacological systems analysis defines EIF4A3 functions in cell-cycle and RNA stress granule formation. Commun Biol 2:1–15. 10.1038/s42003-019-0391-9

52. Egberts J-H, Cloosters V, Noack A, et al (2008) Anti-tumor necrosis factor therapy inhibits pancreatic tumor growth and metastasis. Cancer Res 68:1443–1450. 10.1158/0008-5472.CAN-07-5704

53. Tu M, Klein L, Espinet E, et al (2021) TNF-α-producing macrophages determine subtype identity and prognosis via AP1 enhancer reprogramming in pancreatic cancer. Nat Cancer 2:1185–1203. 10.1038/s43018-021-00258-w

54. Zhao X, Fan W, Xu Z, et al (2016) Inhibiting tumor necrosis factor-alpha diminishes desmoplasia and inflammation to overcome chemoresistance in pancreatic ductal adenocarcinoma. Oncotarget 7:81110–81122. 10.18632/oncotarget.13212

55. Schlautmann LP, Gehring NH (2020) A Day in the Life of the Exon Junction Complex. Biomolecules 10:866. 10.3390/biom10060866

56. Wang Z, Murigneux V, Le Hir H (2014) Transcriptome-wide modulation of splicing by the exon junction complex. Genome Biol 15:551. 10.1186/s13059-014-0551-7

57. Tang Z, Li C, Kang B, Gao G, Li C, Zhang Z (2017) GEPIA: a web server for cancer and normal gene expression profiling and interactive analyses. Nucleic Acids Res 45:W98–W102. 10.1093/nar/gkx247

