## Supplementary figures and images for "The Exon Junction Complex component EIF4A3 plays a splicing-linked oncogenic role in Pancreatic Ductal Adenocarcinoma"

### Supp Fig 1

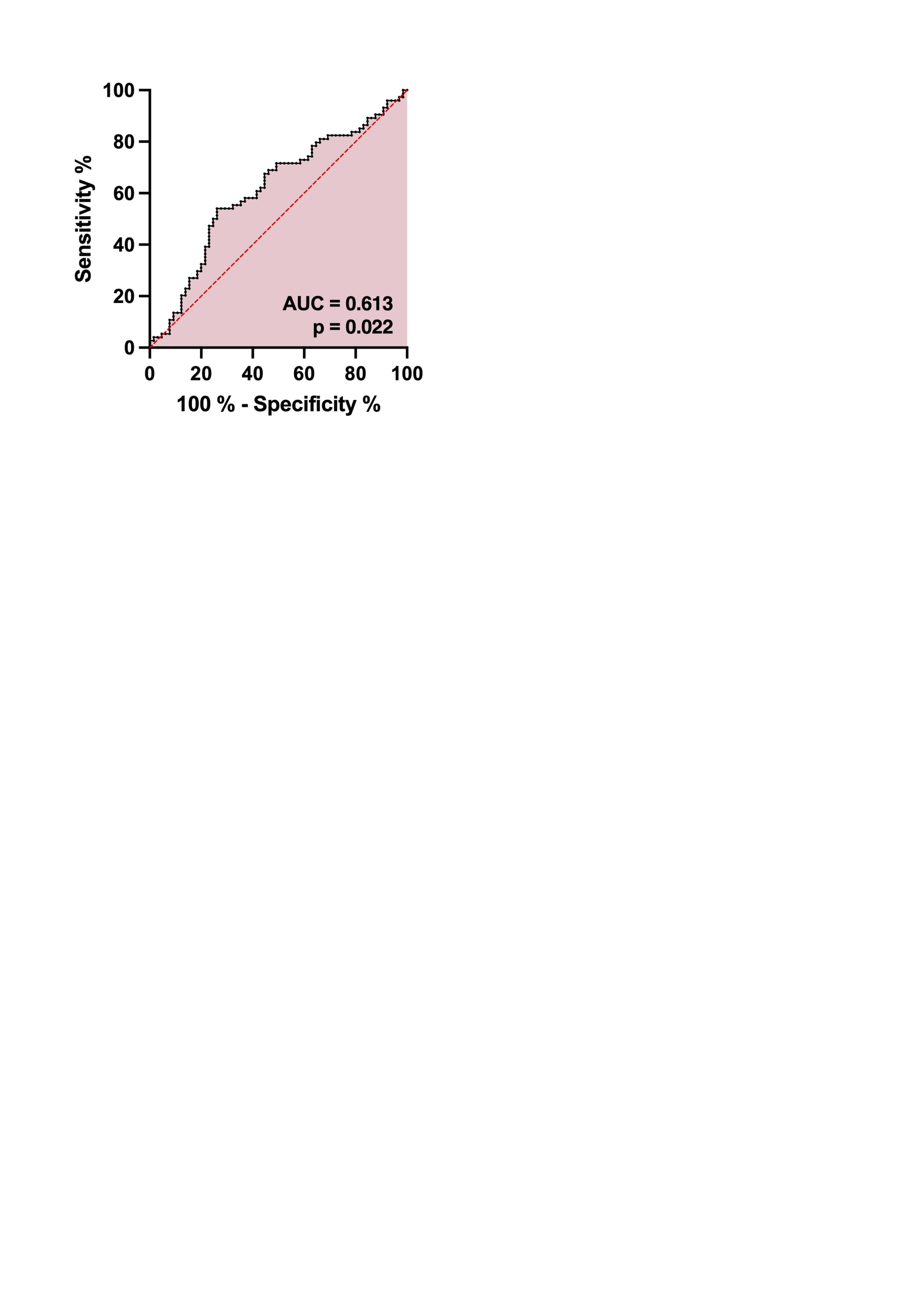

### Supp Fig 2

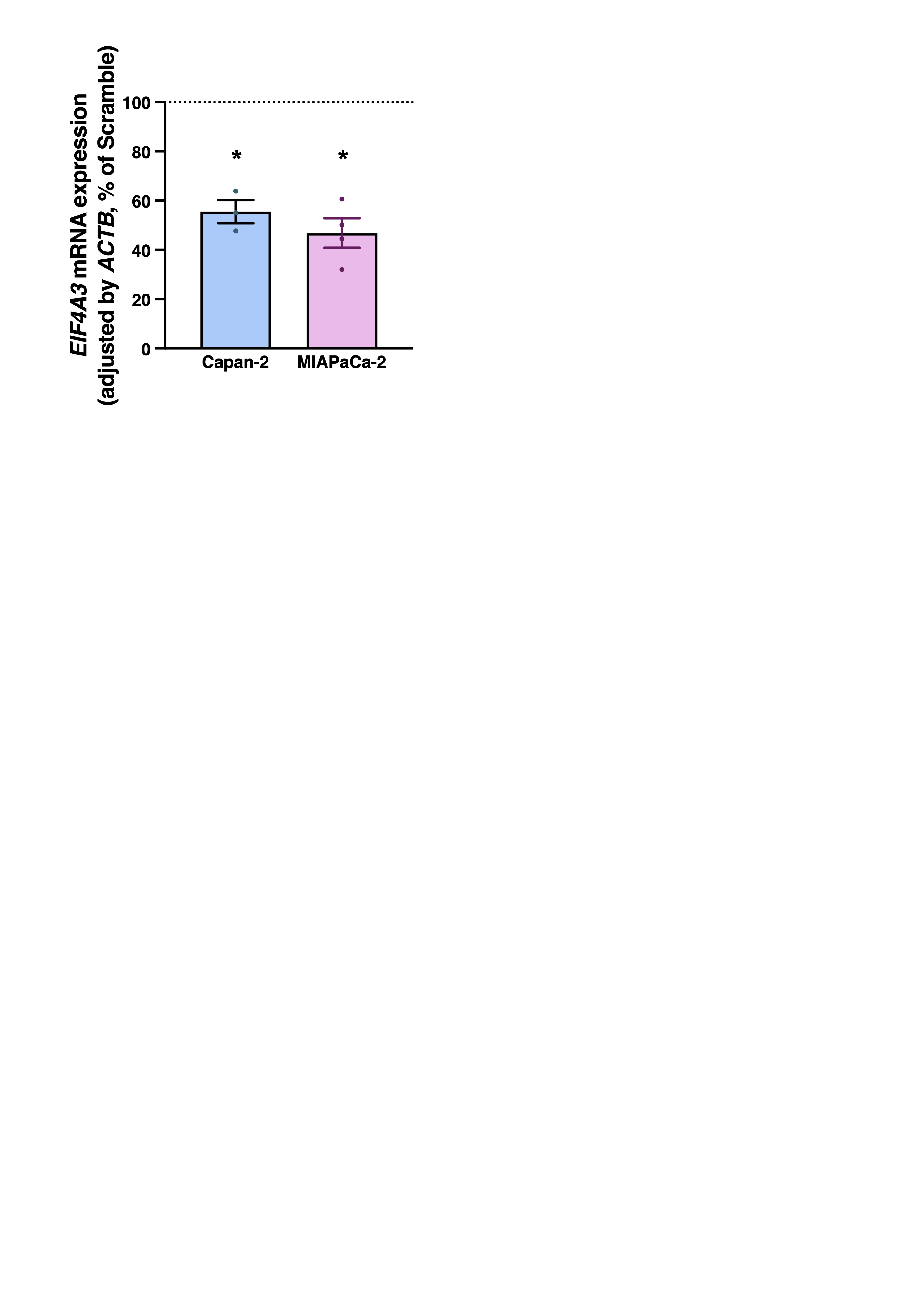
